# Differential contribution of p300 and CBP to regulatory elements in mESCs

**DOI:** 10.1101/806869

**Authors:** Sara Martire, Aishwarya Sundaresan, Laura A. Banaszynski

## Abstract

The transcription coactivators CREB binding protein (CBP) and p300 are highly homologous acetyltransferases that mediate histone 3 lysine 27 acetylation (H3K27ac) at regulatory elements such as enhancers and promoters. Although in most cases, CBP and p300 are considered to be functionally identical, both proteins are indispensable for development and there is evidence of tissue-specific nonredundancy. However, characterization of chromatin and transcription states regulated by each protein is lacking. In this study we analyze the individual contribution of p300 and CBP to the H3K27ac landscape, chromatin accessibility, and transcription in mouse embryonic stem cells (mESC). We demonstrate that p300 is responsible for the majority of H3K27ac in mESCs and that loss of acetylation in p300-depleted mESCs is more pronounced at enhancers compared to promoters. While loss of either CBP or p300 has little effect on the open state of chromatin, we observe that distinct gene sets are transcriptionally dysregulated upon depletion of p300 or CBP. Transcriptional dysregulation is generally correlated with dysregulation of promoter acetylation upon depletion of p300 (but not CBP) and appears to be relatively independent of dysregulated enhancer acetylation. Interestingly, both our transcriptional and genomic analyses demonstrate that targets of the p53 pathway are stabilized upon deletion of p300, suggesting that this pathway is highly prioritized when p300 levels are limited.

## Introduction

Regulatory elements are defined by an open chromatin state and the binding of transcription factors, which in turn contributes to the recruitment of transcription coactivators such as CBP and p300 (Dancy and Cole 2015; Long et al. 2016). In addition to facilitating specific protein interactions at regulatory elements, CBP and p300 acetylate both histone and nonhistone proteins and are specifically responsible for histone acetylation on H3K27 (Zeng et al. 2008; Tie et al. 2009; Pasini et al. 2010; Jin et al. 2011; Dancy and Cole 2015). Once acetylated, these lysines can be recognized by bromodomains, binding motifs that are found in a variety of nuclear proteins associated with transcription (Fujisawa and Filippakopoulos 2017). It then follows that CBP and/or p300 recruitment to regulatory elements, specifically enhancers and promoters, and the subsequent acetylation of these regions is strongly correlated with transcriptional output.

Traditionally and biochemically, CBP and p300 are considered to be a single entity. They share a high degree of sequence identity in structured domains, including their enzymatic HAT domains and their bromodomains (Liu et al. 2008; Zeng et al. 2008). However, CBP and p300 exhibit much lower homology outside of predicted domains and have been reported to interact with different protein partners (Dancy and Cole 2015). A recent study also reports differences in substrate specificity and selectivity based on enzyme levels (Henry et al. 2015). Further, CBP and p300 are each required for mammalian development (Yao et al. 1998; Tanaka et al. 2000) and certain cell types have been shown to be more tolerant of CBP or p300 loss compared to the whole organism (Kasper et al. 2006; Oliveira et al. 2006; Xu et al. 2006; Fauquier et al. 2018), suggesting that CBP and p300 function nonredundantly in vivo.

Previous studies addressing unique functions of p300 and CBP have focused mainly on differences at the level of cellular phenotype corroborated by analysis of transcriptional state in the absence of each coactivator (Kasper et al. 2006; Xu et al. 2006; Fang et al. 2014; Fauquier et al. 2018). Although several studies assess differential binding profiles of p300 and CBP in cultured cells (Ramos et al. 2010; Ianculescu et al. 2012; Kasper et al. 2014), the distinct roles of p300 and CBP in establishing functional chromatin states at a genome-wide level has not been directly explored.

In this study, we analyzed the individual contribution of p300 and CBP to the H3K27ac landscape, chromatin accessibility, and transcription in mESC. To study this question, we used isogenic wild-type and p300 knockdown and CBP knockdown stable lines. We performed H3K27ac chromatin immunoprecipitation followed by high throughput sequencing (ChIP-seq) to assess CBP and p300 contribution to acetylation at enhancers and promoters. We next performed the Assay for Transposase Accessible Chromatin with high-throughput sequencing (ATAC-seq) to assess genome-wide changes in DNA accessibility, which provides a direct measure of chromatin remodeling activity by CBP or p300 complexes. Finally, we performed RNA-seq to link changes in chromatin landscape to gene expression. Overall, we identify distinct roles for CBP and p300 in maintaining H3K27ac at specific regions of the genome. Changes in acetylation do not correlate with changes in DNA accessibility, suggesting that once established, open chromatin states are independent of both the charge neutralization and protein scaffolding provided by lysine acetylation. Overall, we find that p300-mediated acetylation of promoters correlates more strongly with transcription than enhancer acetylation, and that the effects of CBP on transcription are independent of its acetyltransferase activity. Finally, when p300 levels are limiting, we observe a strong propensity toward maintaining acetylation of the p53 pathway, suggesting a previously unrecognized role for p53 as a high-value target of p300 in mESCs.

## Results

### p300 Maintains Enhancer Acetylation in mESCs

To understand the individual contributions of p300 and CBP to the overall level of CBP and p300 in mESCs, we first compared expression levels in wild-type mESCs based on RNA-seq. We found that while p300 transcripts are present at higher levels, both p300 and CBP transcripts are above the median expression level found in mESCs (Figure 1A). We then used shRNA to deplete either p300 or CBP from mESCs. These experiments were performed in biological duplicate and using mESCs treated with a scrambled shRNA sequence as control. Treatment with either p300 or CBP shRNA resulted in reduction of both the targeted transcript and its protein (Figure 1B, 1C). While knockdown of CBP had little apparent effect on p300 protein levels, we observed a slight increase in CBP protein levels upon p300 depletion (Figure 1C). Knockdown of either p300 or CBP had little effect on the pluripotent state of the mESCs based on transcript analysis of important regulators of stemness (e.g., Oct4, Nanog) and alkaline phosphatase staining (Suppl. Figure 1A, 1B).

**Figure 1:**
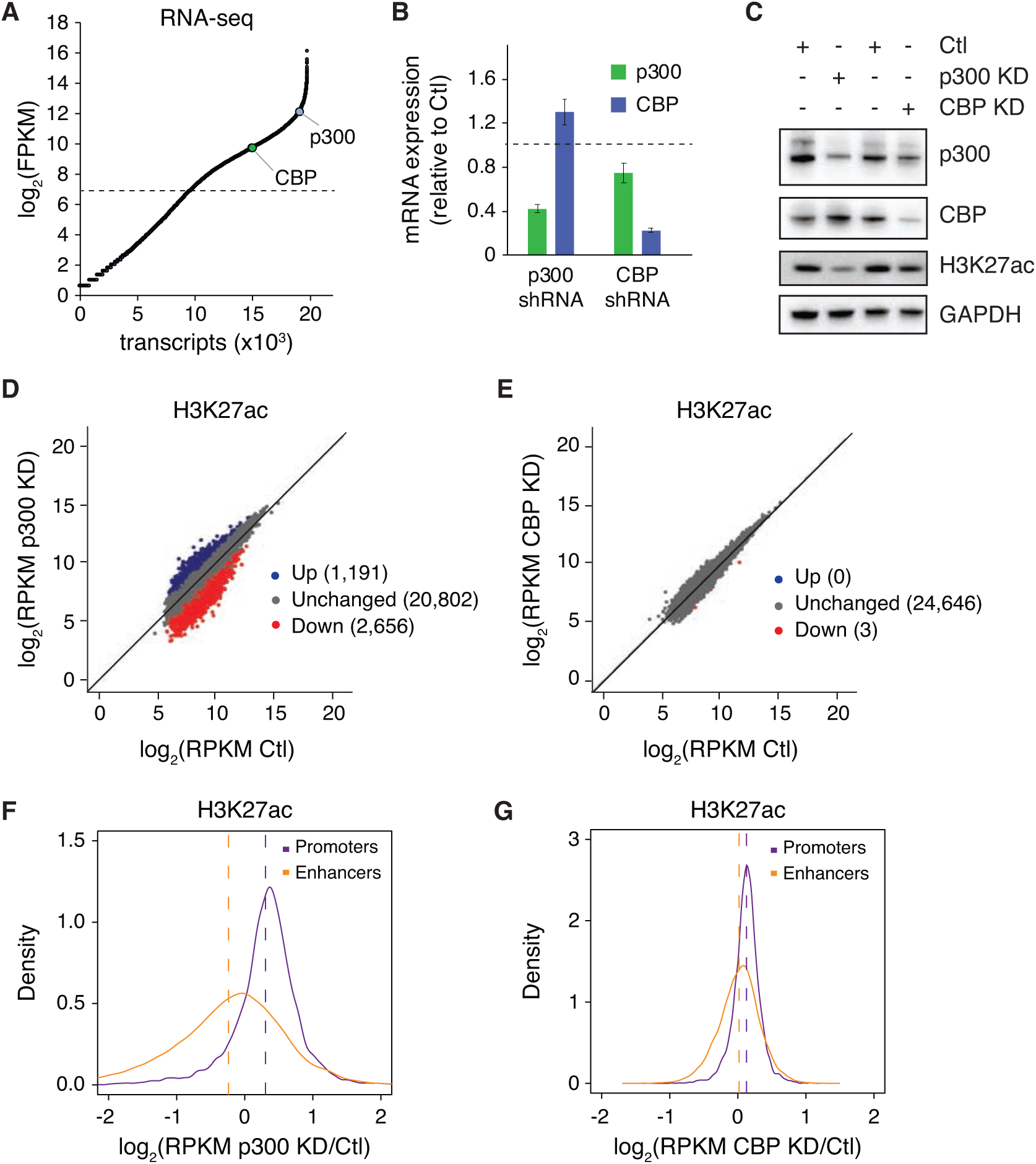
p300 Maintains Enhancer Acetylation in mESCs. (A) Log_2_(FPKM) expression of transcripts in WT mESC cells quantified by RNA-seq of two biological replicates, showing the expression of p300 and CBP. Horizontal dashed line represents the median expression level. (B) Transcript levels (RT-qPCR) of p300 and CBP in mESCs transfected with scramble shRNA or either p300 or CBP shRNA. (C) Immunoblot of whole cell lysates from mESCs transfected with scramble shRNA or either p300 or CBP shRNA. (D,E) H3K27ac ChIP-seq enrichment in mESCs expressing scramble control (Ctl) shRNA compared with (D) p300 shRNA (p300 KD) or (E) CBP shRNA (CBP KD). Blue and red dots represent differential ChIP-seq regions at which read density increased or decreased by 2-fold or more (FDR < 0.05), respectively, in three independent biological replicates. (F,G) Fold-change ratio (log_2_) of H3K27ac enrichment at promoters and enhancers in mESCs expressing scramble control (Ctl) shRNA compared with (F) p300 shRNA (p300 KD) or (G) CBP shRNA (CBP KD). Data to the right of log2(0) = 1 indicate regions with increased H3K27ac after acetyltransferase knockdown, while data to the left indicate regions with reduced H3K27ac.

Although lysines in both histones and non-histone proteins are known to be redundantly modified by both CBP/p300 and other acetyltransferases, H3K27 appears to be exclusively acetylated by CBP/p300. We therefore assessed global levels of H3K27ac upon p300 or CBP knockdown. Interestingly, we observed reduced levels of global H3K27ac after depletion of p300 but not CBP (Figure 1C), indicating that p300 is responsible for the majority of H3K27ac in mESCs. To determine the effect of CBP/p300 loss on H3K27ac genome-wide, we performed chromatin immunoprecipitation followed by sequencing (ChIP-seq) using an H3K27ac antibody.

First, we used triplicate analysis to identify all regions of H3K27ac enrichment in wild-type mESCs treated with control shRNAs (n = 24,649). We then compared H3K27ac enrichment at these regions in mESCs depleted of either p300 or CBP (Figure 1D, 1E), using a 2-fold change cutt-off and statistical significance (p < 0.05) to identify differentially acetylated regions. In agreement with our observation that p300 is responsible for global H3K27ac levels in mESCs (Figure 1C), we found many regions of altered H3K27ac enrichment in p300-depleted mESCs (Figure 1D), whereas H3K27ac enrichment in CBP-depleted mESCs was unaffected and strikingly similar to control mESCs (Figure 1E). While we do observe a number of regions with increased H3K27ac (n = 1,191), the majority of dysregulated regions show reduced H3K27ac upon depletion of p300 under these criteria (n = 2,656).

As H3K27ac is a mark that correlates with both active enhancers and promoters, we next asked how p300 depletion affected these regions individually.Using H3K27ac enrichment data from control mESCs, we defined regions of promoter enrichment to be within ±3 kb of an annotated transcription start site, resulting in 7,336 promoter regions of H3K27ac enrichment. All other regions were annotated as enhancers (n = 16,268). Although both regions showed similar levels of H3K27ac enrichment in control mESCs (Suppl. Figure 1C), only promoters showed high levels of H3K4me3, indicating the validity of this approach (Suppl. Figure 1D). We next compared differences in H3K27ac enrichment at promoters and enhancers in the context of either p300 or CBP depletion (Figure 1F, 1G), plotted as a direct comparison of enrichment at specific loci in acetyltransferase-depleted mESCs versus control mESCs (i.e., fold-change values less than log2(0) = 1 represent regions with reduced acetylation in mESCs depleted of p300 or CBP). Interestingly, in p300-depleted mESCs we observed a clear delineation between enhancers and promoters with enhancers showing a greater loss of H3K27ac enrichment than promoters when p300 levels are limited (Figure 1F). Additionally, in agreement with our global and genomic analyses (Figure 1C, 1E), we observe little change in either promoter or enhancer H3K27ac enrichment after CBP knockdown compared to control mESCs (Figure 1G).

Together, these findings demonstrate that p300, and not CBP, is responsible for the majority of H3K27ac in mESCs. Further, enhancers are more sensitive and show greater reductions in H3K27ac enrichment compared to promoters when p300 levels are limited.

### Chromatin Accessibility Is Independent of CBP/p300 Levels in mESCs

CBP/p300 recruitment and subsequent H3K27ac enrichment are a hallmark of active promoters and enhancers, which can be defined in part by their open chromatin state. Since we observed reduced H3K27ac at enhancers upon p300 depletion, we next asked whether altering the chromatin modification state would influence the “openness” of these regions. We used ATAC-seq to analyze chromatin accessibility at H3K27ac enriched regions in our control shRNA-treated mESCs as well as mESCs depleted of p300 or CBP (Figure 2). Differential regions of ATAC-seq enrichment were identified using a 2-fold cutt-off and statistical significance (p < 0.05). Surprisingly, given the global reduction of H3K27ac, we detected relatively few regions that experienced loss of chromatin accessibility after p300 depletion (Figure 2A, Suppl. Figure 2A). Moreover, regions with reduced accessibility showed little overlap with regions with reduced H3K27ac (Suppl. Figure 2B). Further analysis comparing either enhancer or promoter ATAC-seq signal in control versus p300-depleted mESCs confirmed that these regions generally maintained their open state despite the loss of acetylation observed at enhancers (Figure 2B, Suppl. Figure 2C). In line with CBP having little effect on H3K27ac enrichment, we observed virtually no change in chromatin accessibility upon CBP depletion (Figure 2C, 2D).

**Figure 2:**
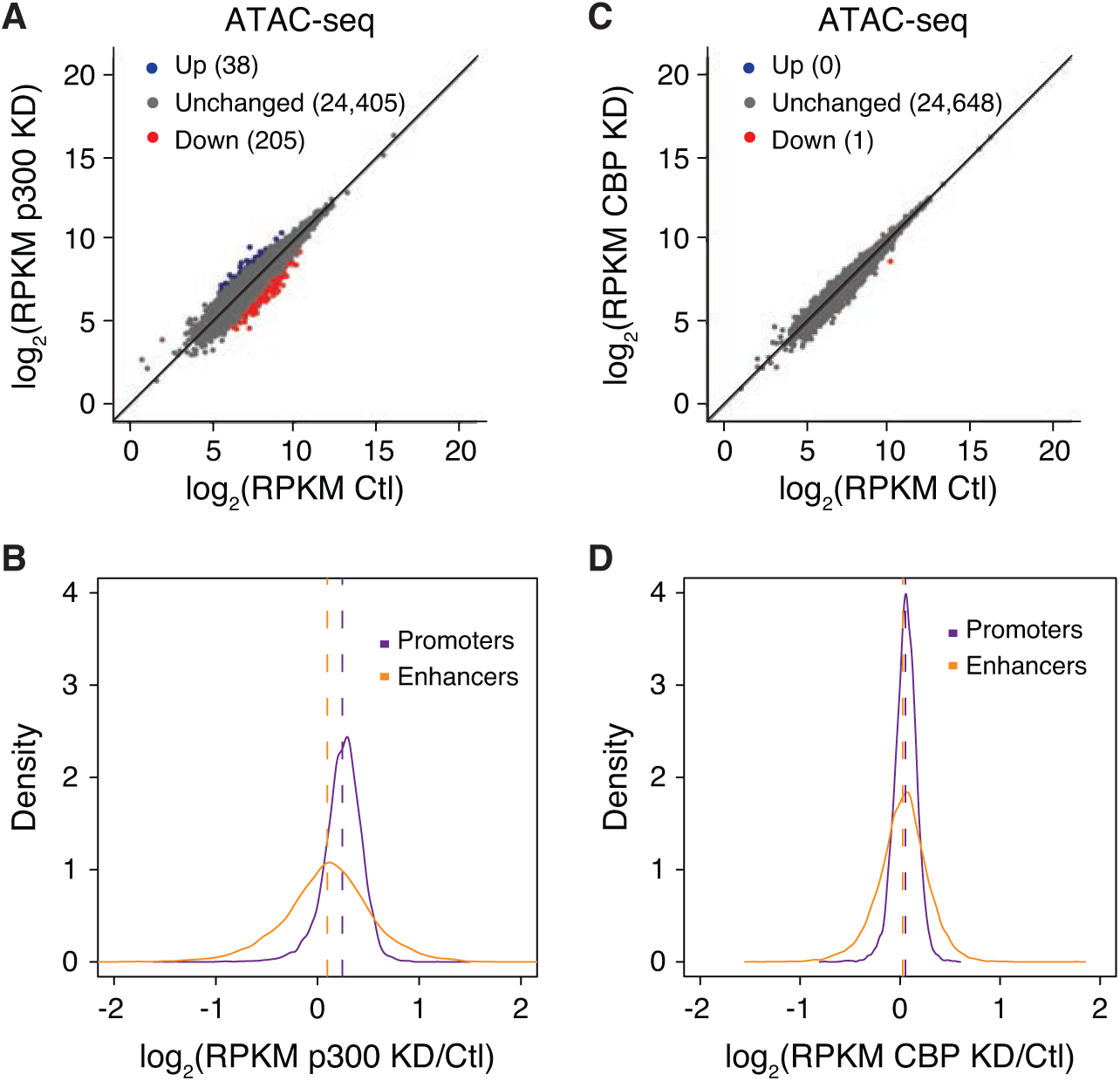
Chromatin Accessibility Is Independent of CBP/p300 Levels in mESCs. (A,C) ATAC-seq levels at H3K27ac-enriched regions in mESCs expressing scramble control (Ctl) shRNA compared with (A) p300 shRNA (p300 KD) or (C) CBP shRNA (CBP KD). Blue and red dots represent regions with differential ATAC-seq signal at which read density increased or decreased by 2-fold or more (FDR < 0.05), respectively, in two independent biological replicates. (B,D) Fold-change ratio (log_2_) of ATAC-seq enrichment at promoters and enhancers in mESCs expressing scramble control (Ctl) shRNA compared with (B) p300 shRNA (p300 KD) or (D) CBP shRNA (CBP KD). Data to the right of log2(0) = 1 indicate regions with increased ATAC-seq signal after acetyltransferase knockdown, while data to the left indicate regions with reduced ATAC-seq signal.

Overall, and in agreement with our previous observations and others (Raisner et al. 2018; Martire et al. 2019), these data demonstrate that maintenance of the open chromatin state in mESCs is independent of the high levels of H3K27ac enrichment observed at these regions in wild-type mESCs.

### Reduced Levels of CBP and p300 Result in Unique Transcriptional Dysregulation

As CBP and p300 are known transcriptional co-activators, we next wanted to determine how loss of p300 or CBP affects the transcriptional state of the cells. We used RNA-seq to compare transcription levels from equal numbers of control and p300- or CBP-depleted mESCs in duplicate using synthetic spike-in standards. Interestingly, the majority of steady-state transcription was unaffected by loss of either p300 or CBP (Figure 3A, 3B). These data are in agreement with previous studies demonstrating that transcription was not uniformly inhibited after deletion of both p300 and CBP from mouse embryonic fibroblasts (Kasper et al. 2010). Regardless, several hundred genes were dysregulated upon reduction of p300 or CBP in mESCs, resulting in both up- and down-regulation of transcript levels (Figure 3A, 3B). Interestingly, we do not observe greater dysregulation of transcription after depletion of p300, the coactivator with a greater effect on H3K27ac.

**Figure 3:**
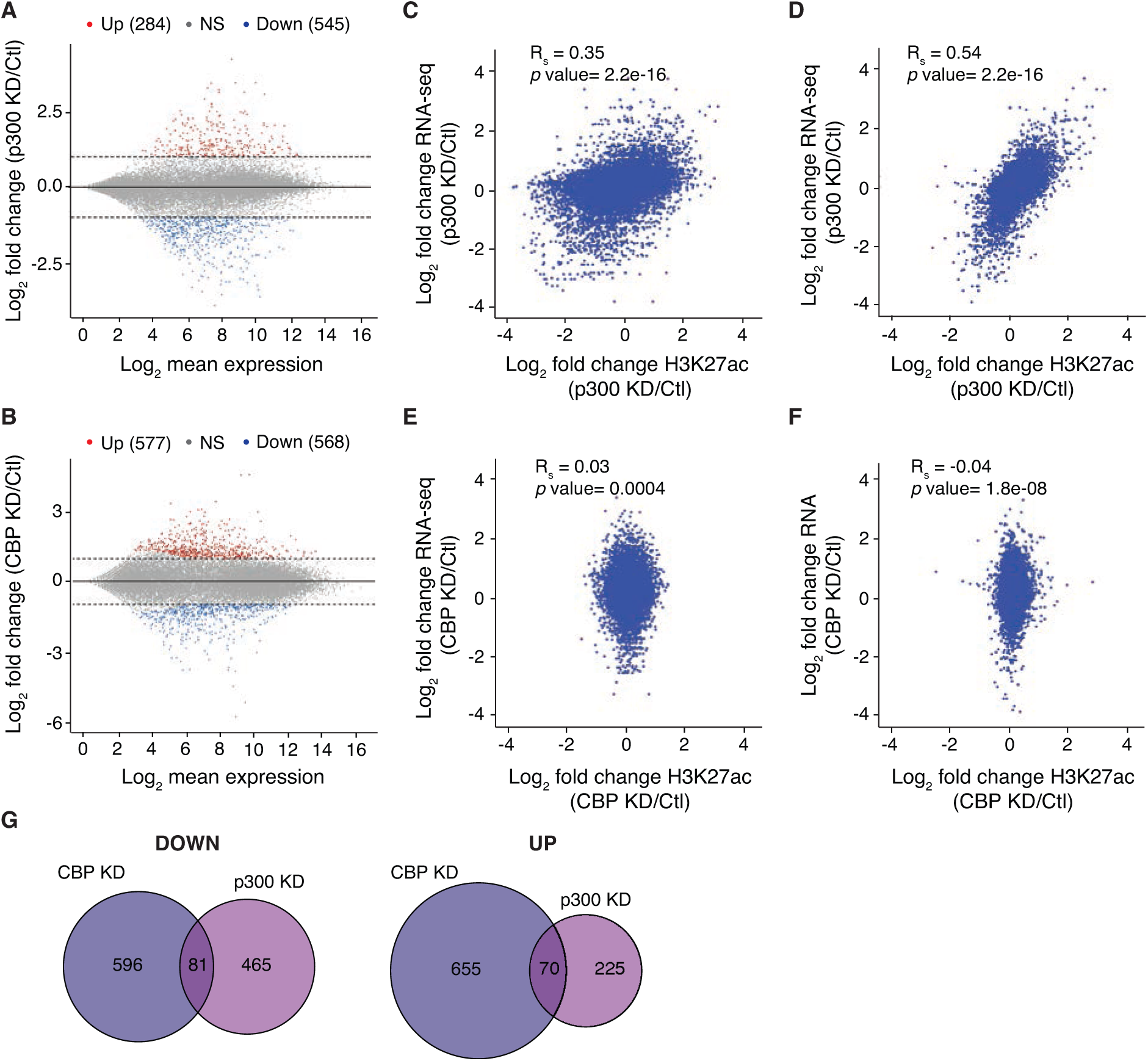
Reduced Levels of CBP and p300 Result in Unique Transcriptional Dysregulation. (A,B) MA plot of gene expression in Ctl and (A) p300 KD and (B) CBP KD mESCs. The x-axis indicates gene counts and the y-axis represents the log_2_(fold-change) in expression for KD versus Ctl mESCs. Dotted lines indicate fold-change >2 and genes in red were differentially expressed (p < 0.05). (C,E) Correlation plot between enhancer H3K27ac ChIP-seq and nearest-neighboring gene RNA-seq for (C) p300 KD and (E) CBP KD represented as log_2_(fold-change) versus Ctl. (D,F) Correlation plot between promoter H3K27ac ChIP-seq and RNA-seq of expressed genes (RPKM >1) for (D) p300 KD and (F) CBP KD represented as log_2_(fold-change) versus Ctl. Spearman correlation tests were performed to determine statistical significance. (G) Venn diagrams showing the relationship between p300 KD and CBP KD RNA-seq in up- and down-regulated genes.

As transcription and acetylation are strongly correlated, we next compared changes in gene expression to changes in genomic acetylation. We focused our analysis in two ways. First, we identified the nearest neighboring genes of enhancers identified in our study and compared changes in expression of these genes to changes in acetylation of their putative enhancers. Second, we compared changes in expression for all expressed genes (RPKM > 1) to changes in acetylation at their promoters. Upon depletion of p300, we observed moderate correlation between changes in transcription and acetylation (Figure 3C, 3D). Notably, p300-dependent changes in acetylation and transcription were more strongly correlated when considering promoter acetylation (Rs = 0.54, Figure 3D), compared to the weak correlation observed between enhancer acetylation and transcription of the nearest neighboring gene (Rs = 0.35, Figure 3C). Given that very little acetylation in mESCs is dependent upon CBP, it is not surprising that we observed no correlation between enhancer acetylation (Rs = 0.03) or promoter acetylation (Rs = −0.04) and changes in transcription upon CBP depletion (Figure 3E, 3F).

Previous studies have suggested that the majority of CBP/p300 function rests with its enzymatic activity, based on similarities observed between enzymatic inhibition of both CBP and p300 and approaches that deplete both CBP and p300 protein levels (Weinert et al. 2018). As loss of either CBP or p300 affects gene expression, but only loss of p300 affects H3K27ac, we asked whether the same sets of genes were dysregulated after CBP or p300 knockdown. Interestingly, we observe very little overlap between genes dysregulated upon CBP or p300 depletion (Figure 3G). Although we cannot discriminate structural from enzymatic functions for p300 in our assay, these data suggest that CBP must regulate gene expression in mESCs through a mechanism that is distinct from its enzymatic function.

Overall, these data suggest that promoter acetylation is a more faithful indicator of transcriptional output than acetylation of nearby regulatory elements in mESCs. Further, the delineation between gene dysregulation upon depletion of p300 and CBP, coupled with CBP’s lack of influence on acetylation, suggests that CBP may play more structural roles in gene regulation in mESCs.

### p53 binding is correlated with maintenance of H3K27ac upon depletion of p300

As we observed that p300 and CBP regulate different sets of genes in mESCs, we performed gene set enrichment analysis (GSEA) of our RNA-seq data to understand the pathways regulated by p300 and CBP, respectively (Figure 4A, Suppl. Figure 3A). Surprisingly, one of the most enriched terms observed after p300 knockdown was upregulation of the p53 pathway (Figure 4A). This pathway was not observed in GSEA analysis of CBP-regulated genes (Suppl. Figure 3A). We further confirmed that p300 depletion led to dysregulation of several p53 targets (Figure 4B), demonstrating that p53-activated genes are upregulated upon p300 depletion and, likewise, that negatively regulated p53-targets are downregulated. Although p53 is a known target of p300 (Gu and Roeder 1997; Sakaguchi et al. 1998; Liu et al. 1999) we did not observe decreased p53 acetylation upon p300 depletion (Suppl. Figure 3B). Further, despite transcriptional dysregulation of the p53 pathway, we did not observe clear signs of baseline DNA damage or sensitivity to irradiation after depletion of p300 in mESCs (Suppl. Figure 3C).

**Figure 4:**
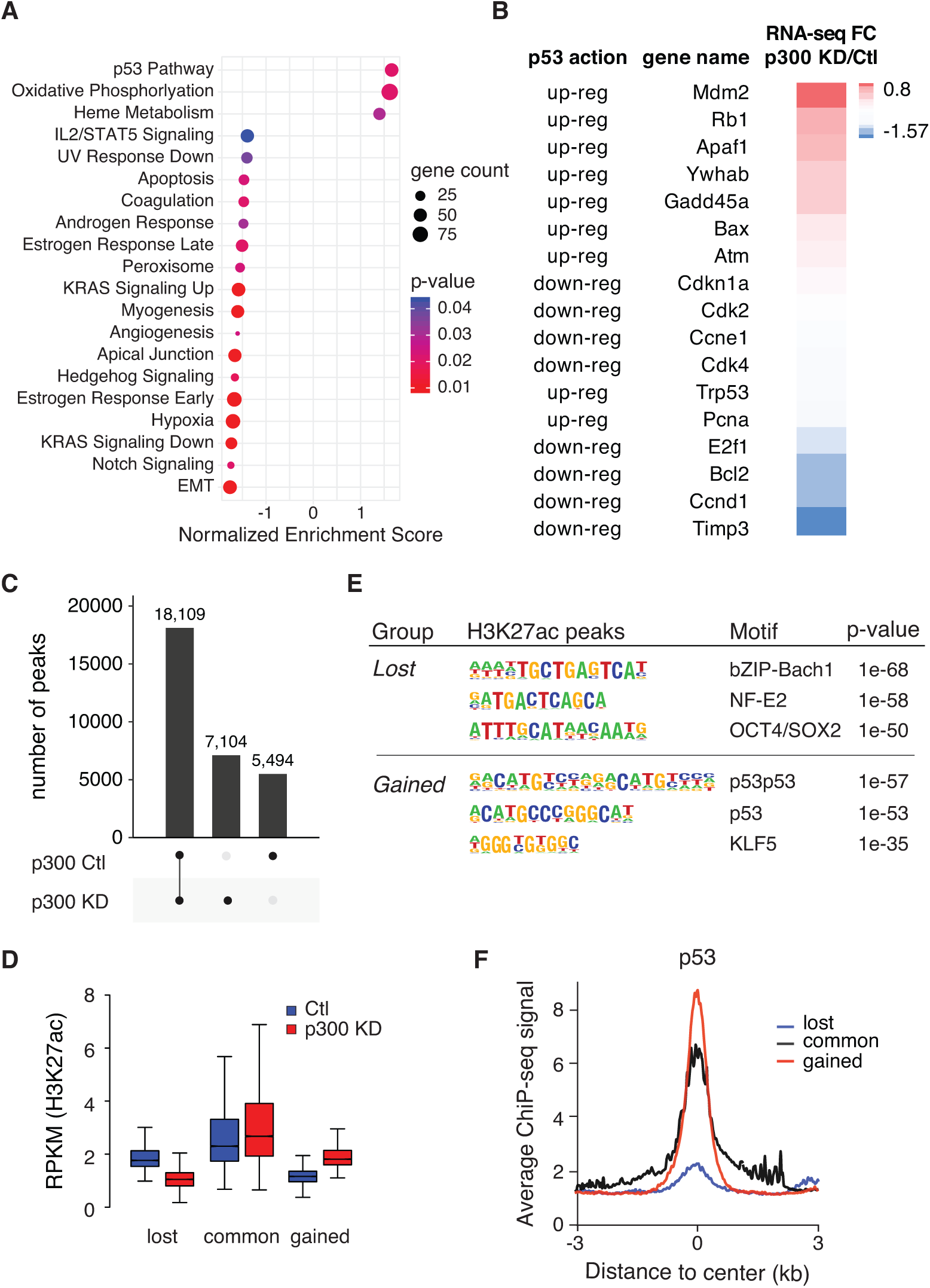
The p53 Pathway Remains Highly Acetylated when p300 Levels are Limited. (A) GSEA pathway analysis of significantly regulated genes (based on RNA-seq) in p300 KD mESC compared to Ctl mESC. The normalized enrichment score (NES) is indicated for each gene set. (B) Fold-change RPKM (RNA-seq) of p300 KD versus Ctl for genes belonging to the p53 pathway. Effect of p53 on gene expression is indicated. (C) UpSet representation of regions of H3K27ac enrichment and overlap in Ctl and p300 KD mESCs. (D) Boxplot showing H3K27ac enrichment at lost, common and gained peaks after p300 KD. p < 2.2 x 10^-16^ for all comparisons by Wilcoxon rank sum test. The bottom and top of the boxes correspond to the 25^th^ and 75^th^ percentiles, and the internal band is the 50^th^ percentile (median). The plot whiskers correspond to 1.5× interquartile range and outliers are excluded. (E) Motif enrichment analysis of lost and gained regions of H3K27ac enrichment after p300 KD. Logo and p-values are indicated for each motif. (F) Average profile of p53 ChIP-seq in WT mESCs at lost, common and gained regions of H3K27ac enrichment after p300 KD.

To corroborate our transcriptional analysis, we returned our focus to H3K27ac enrichment profiles in control and p300-depleted mESCs. Using peak calling tools, we identified regions of H3K27ac enrichment that were identified only in our control mESCs (and therefore lost after p300 depletion, referred to as lost), regions that were common to both the control and the p300-depleted mESCs (common), and regions that were found only in p300-depleted mESCS (gained) (Figure 4C). We validated our peak calling by demonstrating the expected changes in H3K27ac enrichment at these regions in control and p300-depleted mESCs (Figure 4D).

To determine regulatory networks underlying H3K27ac enrichment in control and p300-depleted mESCs, we used HOMER (Heinz et al. 2010) to predict transcription factor binding motifs enriched at these regions. In regions at which H3K27ac is lost upon p300 depletion, one of the most enriched motifs was Oct4/Sox2 (p = 1 x 10^-50^), which is highly correlated with pluripotency (Figure 4E). Likewise, these regions are highly enriched with the pluripotency transcription factors, Oct4, Sox2, and Nanog(Chronis et al. 2017), in wild-type mESCs (Suppl. Figure 3D). Interestingly, we found that regions with increased H3K27ac (gained) upon p300 depletion were enriched with the p53 binding motif (Figure 4E). Finally, we performed p53 ChIP-seq in wild-type mESCs to determine p53 enrichment across these regions. We found that p53 was greatly enriched in the regions that maintain or gain acetylation upon p300 depletion compared to regions that depend upon p300 to maintain high levels of H3K27ac (Figure 4F). Further, pluripotency transcription factors showed relatively low enrichment at regions that gain H3K27ac after p300 depletion (Suppl. Figure 3D). Interestingly, and in line with previous studies suggesting pioneer factor activity for p53 (Sammons et al. 2015; Younger and Rinn 2017), we observe that the “gained” p53-bound regions exist in a more closed state compared with the regions highly bound by pluripotency transcription factors in control mESCs (Suppl. Figure 3E).

Overall, these data demonstrate that H3K27ac of genes involved in the p53 regulatory pathway remains high when p300 is limited, suggesting that the p53 pathway is highly prioritized amongst targets of p300 in mESCs. Further, regions that lose H3K27ac upon p300 depletion exist in a more open chromatin state and are highly bound by pluripotency transcription factors in control mESCs, suggesting that a robust gene regulatory network is able to maintain chromatin accessibility and gene expression in mESCs when p300 activity is reduced.

## Discussion

Based on their high degree of sequence identity and assessment of activity in vitro, CBP and p300 are thought to function redundantly and they are often referred to as a single entity. However, several pieces of evidence support a model in which p300 and CBP are uniquely required for context-dependent gene expression. Most notably, homozygous null mutations of either coactivator result in early embryonic lethality (Yao et al. 1998; Tanaka et al. 2000). Further, conditional deletion or depletion of p300 or CBP suggests selective roles for these two proteins in a number of adult tissues and contexts (Kasper et al. 2006; Oliveira et al. 2006; Xu et al. 2006; Fauquier et al. 2018). In line with these observations, our analysis demonstrates that p300 and CBP carry out unique functions in embryonic stem cells. Specifically, we find that H3K27ac levels in mESCs are generally independent of CBP and that p300 is the acetyltransferase responsible for the majority of H3K27ac in mESCs.

Given the high correlation between the “open” chromatin state and histone acetylation, we were surprised that reduced p300 and the subsequent reduction in H3K27 acetylation did not affect chromatin accessibility at acetylated regions. Nevertheless, this data aligns with our previous observation that a significant reduction of p300 HAT activity (i.e., H3K27ac) fails to “close” chromatin at p300 targets (Martire et al. 2019). It is also supported by a recent study in which small-molecule inhibition of the CBP/p300 bromodomain reduced global H3K27ac without changing the open chromatin state (Raisner et al. 2018). Taken together, these data suggest that neither the HAT activity of p300 nor more fundamentally its presence is required to maintain open chromatin at regulatory regions. Interestingly, we find that the regions that lose the most H3K27ac when p300 levels are limited are also regions that are highly bound by pluripotency-specific transcription factors. This observation is in line with previous data suggesting that genes with greater local concentration and diversity of coactivators are less reliant on CBP/p300 to maintain steady state gene expression(Kasper et al. 2010).

Although acetylation has long been correlated with gene expression, we and others find that reduced acetylation is well-tolerated in mESCs (Dorighi et al. 2017; Rickels et al. 2017; Martire et al. 2019). It is possible that under culture conditions regulatory elements are acetylated more robustly than is required for a functional readout of this modification. Such “buffering” could explain why a global loss in H3K27ac levels has modest effect on steady state transcription. In agreement, two recent studies reported that reduced MLL activity at enhancers resulted in reduced H3K27ac at these regions with little effect on global transcription (Dorighi et al. 2017; Rickels et al. 2017). Nevertheless, mutations in the enzymatic HAT domain of p300 and CBP are highly recurrent in many tumor types, suggesting important context-dependent roles for H3K27ac.

Although p300 and CBP function is described at both promoters and enhancers(Dancy and Cole 2015) our findings indicate that enhancers are the principal sites affected by reduced levels of p300. This observation is in agreement with patterns of H3K27ac loss after treatment with CBP/p300 bromodomain inhibitor (Raisner et al. 2018). Previous studies have interpreted results such as these as an indication that CBP/p300 play a more important role at enhancers. In contrast, we suggest a new vision, that when p300 levels are limited, loss of enhancer acetylation is tolerated while maintenance of promoter acetylation is more important to safeguard critical cell functions. In support, we observe stronger correlation between changes in promoter acetylation and transcription than changes in enhancer acetylation and transcription of the nearest neighboring genes upon depletion of p300.

Intriguingly, regions that maintain or gain acetylation when p300 is limited are enriched with p53 binding motifs and are indeed highly bound by p53 in wild-type mESCs. In agreement with previous studies (Sammons et al. 2015; Younger and Rinn 2017), we find that p53-bound enhancers are located within regions of relatively less accessible chromatin, suggesting that p53 may regulate enhancer activity, in part, by modulating chromatin accessibility given the appropriate cellular context. Previous studies demonstrate that, in line with its function as a tumor suppressor, p53 activity is low in mESCs and increases to promote differentiation in part by silencing critical pluripotency factors (Lin et al. 2005). In fact, p53 knockdown has been used as a tool to promote cellular reprogramming towards the iPS state (Hong et al. 2009; Kawamura et al. 2009; Marion et al. 2009; Utikal et al. 2009). However, we see no indication of differentiation after p300 knockdown. It is possible that the maintenance of p53-associated acetylation in p300-depleted cells is to ensure proper mESC differentiation. In support, findings from numerous in vivo and in vitro studies now suggest that p53 plays a gatekeeper function to ensure high-fidelity development, promoting specification of cells that undergo cell-cycle arrest and are apoptosis-competent (Aylon and Oren 2016). This connection is highly deserving of future studies.

In sum, we demonstrate unique functions for the acetyltransferases p300 and CBP in mESC. We find that mESCs are quite tolerant of reduced H3K27ac at enhancers, and that promoter acetylation is a stronger indicator of transcriptional competence. Further, we identify the p53 pathway as an important target of p300 in mESCs. Taken together, our study offers new insights into an important family of transcriptional coactivators.

## Methods

### mESC Culture

Mouse embryonic stem cell lines (mESCs) were cultured on gelatin-coated plates under standard serum/LIF conditions at 37 °C with 5% CO_2_ (KO-DMEM, 2 mM Glutamax, 15% ES grade fetal bovine serum, 0.1 mM 2-mercaptoethanol, 1x Pen/Strep, 1x NEAA and leukemia inhibitory factor (LIF)). During thawing and early passages, cells were maintained on an irradiated feeder layer. To remove feeders, cells were passaged at least two passages off of feeders onto gelatin-coated plates. mESCs were routinely tested for mycoplasma.

### shRNA transduction

For p300 and CBP knockdown, 5 μg of plasmids (Dharmacon) were packaged with 5 μg psPAX2, and 0.5 μg VSVG plasmids and transfected in serum-free media into 3×10^6^ 293T cells in a 10 cm^2^ tissue culture dish using Lipofectamine 3000. Lentivirus-containing supernatants were harvested 48 and 72 hrs post-transfection, pooled, and concentrated 10x with Lenti-X (Clontech, 63123). 2×10^5^ WT mESCs were incubated with 0.2 ml concentrated lentivirus and polybrene (8 μg/ml). The next day, the media was replaced with complete mESC culture media containing 1 μg/ml puromycin (for p300 plasmid) or 400 ug/ul (for CBP plasmid) of G418. After 4 days of selection, mESCs were used for downstream analysis.

### Antibodies

H3K27ac (39133, Active Motif. Lot # 31814008), Spike-In antibody (61686, Active Motif, Lot# 00419007), p300 (sc-584, Santa Cruz, Lot # F3016), Gapdh (2118, Cell Signaling, Lot # 10), CBP(D6C5) (7389S, Cell Signaling), p53(CM5) (NCL-L-p53-CM5p, Leica Biosystems), PKCs p2056 (ab18192, abcam), KAP1/TRIM29 pS824 (A300-767A, Bethyl), Acetyl-p53 (Lys379) (2570, Cell Signaling)

### Irradiation of cells

Gamma radiation experiments were conducted at UT Southwestern Medical Center using a 137Cs source irradiator. mESC cells were irradiated with 10 Gy of irradiation (IR) for 3 minutes and allowed to recover for 30 min.

### Chromatin Immunoprecipitation (ChIP)

Native and crosslinking (X) ChIP were performed with 5×10^6^ and 1×10^8^ cells, respectively, as previously described (Martire et al. 2019) with slight modifications. A spike-in normalization strategy was used to normalize all ChIP-seq data to reduce the effects of technical variation and sample processing bias. Spike-In chromatin (Active Motif, 53083) and Spike-in antibody (Active Motif, 61686) were used according to manufacturer instructions.

#### Native ChIP

Cells were trypsinized, washed and subjected to hypotonic lysis (50 mM TrisHCl pH 7.4, 1 mM CaCl_2_, 0.2% Triton X-100, 10 mM NaButyrate, and protease inhibitor cocktail (Roche)) with micrococcal nuclease for 5 min at 37 °C to recover mono- to tri-nucleosomes. Nuclei were lysed by brief sonication and dialyzed into RIPA buffer (10 mM Tris pH 7.6, 1 mM EDTA, 0.1% SDS, 0.1% Na-Deoxycholate, 1% Triton X-100) for 2 hr at 4 °C. Soluble material was incubated with 3-5 μg of antibody bound to 50 μl protein A or protein G Dynabeads (Invitrogen) and incubated overnight at 4 °C, with 5% reserved as input DNA. Magnetic beads were washed as follows: 3x RIPA buffer, 2x RIPA buffer + 300 mM NaCl, 2x LiCl buffer (250 mM LiCl, 0.5% NP-40, 0.5% NaDeoxycholate), 1x TE + 50 mM NaCl. Chromatin was eluted and treated with RNaseA and Proteinase K. ChIP DNA was purified and dissolved in H_2_O.

#### X-ChIP

Cells were crosslinked with 1% formaldehyde for 10 min at room temperature and quenched with 0.125 M glycine. Cells were divided in 3 batches and ChIPs were perfomed in parallel and then pool together. Cells were resuspended in Farnham Lysis Buffer (5 mM PIPES pH 8, 85 mM KCl, 0.5% NP-40, 1mM DTT, protease and phosphatase inhibitors) and isolated nuclei in Lysis Buffer (1% SDS, 10 mM EDTA, 50 mM Tris (pH7.9), 1 mM DTT, protease and phosphatase inhibitors) and chromatin was sonicated to an average size of 0.3-0.7 kb using Covaris M220 Focused-ultrasonicator. Soluble material was diluted in 10X dilution buffer (0.5% Triton X-100, 2 mM EDTA, 20 mM Tris (pH 7.9), 150 mM NaCl, 1 mM DTT, protease and phosphatase inhibitors) and incubated with 40 μl of antibody bound to 150 μl protein A/G mixed Dynabeads (Invitrogen) and incubated overnight at 4 °C, with 5% reserved as input DNA. Magnetic beads were washed as follows: 1x Low Salt buffer (10 mM TrisHCl pH 8, 2 mM EDTA, 0.1% SDS, 1% Triton X-100, 150 mM NaCl), 1x High Salt buffer (10 mM TrisHCl pH 8, 2 mM EDTA, 0.1% SDS, 1% Triton X-100, 500 mM NaCl), 1x LiCl buffer (10 mM TrisHCl pH 8, 1 mM EDTA, 1% NP-40, 1% NaDeoxycholate, 250 mM LiCl), 1x TE + 50 mM NaCl. Chromatin was eluted and treated with RNaseA and Proteinase K. ChIP DNA was purified and dissolved in H_2_O.

### ChIP-seq

#### ChIP-seq Library Preparation

ChIP-seq libraries were prepared from 5 ng ChIP DNA following the Illumina TruSeq protocol. The quality of the libraries was assessed using a D1000 ScreenTape on a 2200 TapeStation (Agilent) and quantified using a Qubit dsDNA HS Assay Kit (Thermo Fisher). Libraries with unique adaptor barcodes were multiplexed and sequenced on an Illumina NextSeq 500 (paired-end, 33 base pair reads). Typical sequencing depth was at least 30 million reads per sample.

#### ChIP-seq Data Quality Control, Alignment and spike-in normalization

Quality of ChIP-seq datasets was assessed using the FastQC tool. ChIP-seq raw reads were aligned separately to the mouse reference genome (mm10) and the spike-in drosophila reference genome (dm3) using BWA (Li and Durbin 2009). Only one alignment is reported for each read (either the single best alignment or, if more than one equivalent best alignment was found, one of those matches selected randomly). Duplicate reads were filtered using the Picard MarkDuplicates. Uniquely mapped drosophila reads were counted in the sample containing the least number of drosophila mapped reads and used to generate a normalization factor for random downsampling. Reads were converted into bigWig files using BEDTools (v2.29.0) (Quinlan and Hall 2010) for visualization in Integrative Genomics Viewer.

#### Downstream ChIP-seq analysis

- *Peak Calling*. Peak calling was performed using MACS14 (Zhang et al. 2008) software using input as a control in each replicated sample. HOMER mergePeaks (Heinz et al. 2010) was used to get unique peaks from replicates to reduce false positives and retain only robust peaks for further analyses.
- *Average Profiles.* Bigwig files were used to generate average ChIP-seq profiles using deepTools.
- *Heatmaps.* The read densities surrounding 6 kb (± 3 kb) of the peak center of WT H3K27ac peaks were determined and visualized as heatmaps using deepTools (Ramirez et al. 2016).
- *Box plots.* Box plot representations were used to quantitatively assess the read distribution in a fixed window. Box plots are defined by the median, box limits at upper and lower quartiles of 75% and 25%, and whiskers at 90% and 10%. The read distribution surrounding the peak center was calculated and plotted using custom R scripts. Wilcoxon rank sum tests were performed to determine the statistical significance of all comparisons.
- *Differential Expression Analysis (DE) of H3K27ac ChIP-seq.* Differential peaks were found using multiBamSummary (DeepTools) (Ramirez et al. 2016) in a BED-file mode, by using WT H3K27ac peaks as input BED file. The output of multiBamSummary is a compressed numpy array (.npz) that was directly used by the program DE-Seq2 with fold change ≥2 or ≤−2, FDR < 0.05 to calculate and visualize pairwise correlation values between the read coverages using custom R scripts.
- *Density plots.* Density plots representing fold-change differences between samples were generated using custom R scripts. k-s tests were performed to determine the statistical significance of all comparisons.
- *UpSet Plots*. To check the extent of overlap of H3K27ac peaks between samples, UpSet Plots were generated using custom R scripts.
- *Motif analysis.* HOMER findMotifs (Heinz et al. 2010) was used to perform motif analysis on H3K27ac peaks.

### ATAC-seq

ATAC-seq was performed as previously described (Buenrostro et al. 2013) with minor changes. For each sample, 100,000 cells were harvested, washed and lysed with ATAC buffer (Tris 10 mM pH 7.4, 10 mM NaCl, 3 mM MgCl_2_, NP-40 0.1%, Tween-20 0.1%, Digitonin 0.01%). Nuclei were collected and subject to tagmentation at 37 °C for 30 minutes in adjusted tagmentation buffer (2x TD Tagment buffer + Digitonin 0.01% + 5 ul of TDE Tagment DNA enzyme from Illumina). Reaction was stopped with 0.2% SDS and DNA was collected using Qiaquick PCR purification columns and eluted in 10 μl 10 mM Tris, pH 8. Eluted DNA was amplified using NebNext Q5 MM kit and purified using AMPure XP beads (negative and positive selection). Samples were pooled for multiplexing and sequenced using paired-end sequencing on the Illumina NextSeq 500.

#### ATAC-seq Data Quality Control, Alignment and normalization

Quality of the ATAC-seq datasets was assessed using the FastQC tool. The ATAC-seq reads were then aligned to the mouse reference genome (mm10) using BWA (Li and Durbin 2009). For unique alignments, duplicate reads were filtered out. The resulting uniquely mapped reads were normalized to the same read depth across all samples and converted into bigWig files using BEDTools (Quinlan and Hall 2010) for visualization in Integrative Genomics Viewer(Robinson et al. 2011). Heatmaps were generated using deepTools.

#### Downstream ATAC-seq analysis

- *Differential Expression Analysis (DE) of ATAC-seq.* Differential peaks were found using multiBamSummary(DeepTools) (Ramirez et al. 2016) in a BED-file mode, by using WT H3K27ac peaks as BED file. The output of multiBamSummary is a compressed numpy array (.npz) that was directly used by the program DE-Seq2 with fold change ≥2 or ≤−2, FDR < 0.05 to calculate and visualize pairwise correlation values between the read coverages using custom R scripts.
- *Density plots.* Density plots representing fold-change differences between samples were generated using custom R scripts. k-s tests were performed to determine the statistical significance of all comparisons.

### Quantitative RT-PCR and mRNA-seq

mRNA was isolated using QIAGEN RNeasy. 500 ng of total RNA was reverse transcribed using random hexamers and MultiScribe reverse transcriptase. mRNA expression was analyzed by quantitative PCR (qPCR) with SYBR Green using a LightCycler 480 (Roche). All qPCR primer sequences used in this study are listed in Table S1.

For RNA-seq, total mRNA from equal cell numbers was mixed with synthetic RNA standards (ERCC RNA Spike-In Mix, Thermo Fisher) (Jiang et al. 2011). Libraries were prepared according to the Illumina TruSeq protocol and sequenced on an Illumina NextSeq 500 (paired-end, 33 base pair reads).

### Analysis of RNA-seq Data

#### Data Quality Control, Alignment and normalization

Quality of the RNA-seq raw reads was assessed using the FastQC tool. The reads were then aligned to the mouse reference genome (mm10) and the spike-in control ERCC92 using STAR (Dobin et al. 2013). Reads mapping to ERCC92 were counted using htseq-count (Anders et al. 2015) and used to normalize the counts to genes. After normalization, the reads were converted into bigWig files using BEDTools for visualization in Integrative Genomics Viewer (Robinson et al. 2011) or the UCSC genome browser.

#### Downstream RNA-seq analysis

- *Differential Expression Analysis (DE)*. Gene expression level measured as FPKM was determined by the maximum likelihood estimation method implemented in the htseq-count software package with annotated transcripts as references. Differential expression was analyzed using the Student’s t test in the program DE-Seq2 with *p* values corrected for multiple testing.
- *MA Plots.* MA plots were used to graphically represent genes that were upregulated or downregulated by more than 2-fold. The log2 fold change (KO/WT) was plotted on the y-axis versus the log2 mean of normalized counts on the x-axis.
- *Venn Diagrams*. To check the extent of overlap of the Up and Down regulated genes from RNA-seq from p300 and CBP datasets, venn diagrams were generated using custom R scripts.
- *Correlation plots.* Correlation of fold-change differences between samples, comparing H3K27ac levels and RNA expression was generated using custom R scripts. Spearman correlation tests were performed to determine the statistical significance of all comparisons.

### Quantification and Statistical Analysis

To check the significance of all comparisons, Wilcoxon rank sum test was used to calculate p-values for data used to generate boxplots. Two-sample Kolmogorov-Smirnov test was used to calculate p-values to show significant changes between two density curves. Spearman correlation tests were performed to determine the statistical significance of correlation plots.

### Code Availability

Codes to generate figures are available upon request.

### Data Availability

Datasets are deposited in the NCBI Gene Expression Omnibus using the following accession numbers: XXX.

## Acknowledgments

We thank members of the Banaszynski laboratory for helpful discussions; E. Duncan and R. O’Hara for critical comments on this manuscript; A. Davis for experimental assistance with DNA damage assays; UTSW BioHPC for computational infrastructure; UTSW McDermott Center for providing next-generation sequencing services. L.A.B. is a Virginia Murchison Linthicum Scholar in Medical Research (UTSW Endowed Scholars Program) and a Peterson Investigator of the Neuroendocrine Research Foundation (NETRF). This work was supported in part by CPRIT RR140042, The Welch Foundation I-1892, DoD KCRP KC170230, and NIH R35 GM124958 (L.A.B.), the American-Italian Cancer Foundation (S.M.) and the Green Center for Reproductive Biology Sciences.

## Author Contributions

S.M. and L.A.B. conceived and designed the study; S.M. performed experiments and computational analyses; A.S. performed GSEA analysis and assisted with R script generation; L.A.B. supervised the project; S.M. and L.A.B. wrote the manuscript.

---

**Suppl Figure 1:**
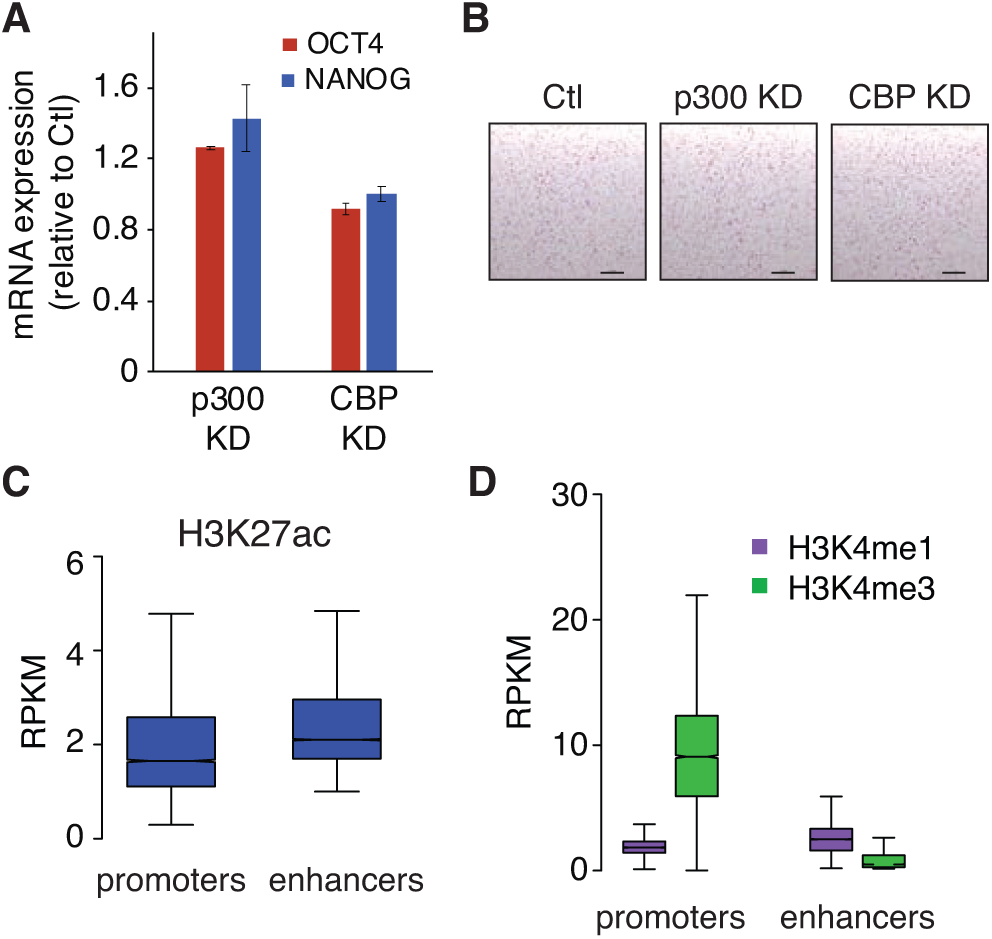
p300 Maintains Enhancer Acetylation in mESCs. (A) Transcript levels (RT-qPCR) of Oct4 and Nanog in p300 and CBP KD mESCs compared to Ctl. (B) Alkaline phosphatase staining of Ctl, p300 KD, and CBP KD mESCs in S/L media. Scale bar = 1 mm. (C) Boxplot showing H3K27ac enrichment at enhancers (n = 16,268) and promoters (n = 7,336) in wild-type cells. p < 2.2 x 10^-16^ for all comparisons by Wilcoxon rank sum test. The bottom and top of the boxes correspond to the 25^th^ and 75^th^ percentiles, and the internal band is the 50^th^ percentile (median). The plot whiskers correspond to 1.5× interquartile range and outliers are excluded. (D) Boxplot showing H3K4me1 and H3K4me3 enrichment at enhancers (n = 16,268) and promoters (n = 7,336) in wild-type cells. p < 2.2 x 10^-16^ for all comparisons by Wilcoxon rank sum test. Boxplots displayed as in panel C.

**Suppl Figure 2:**
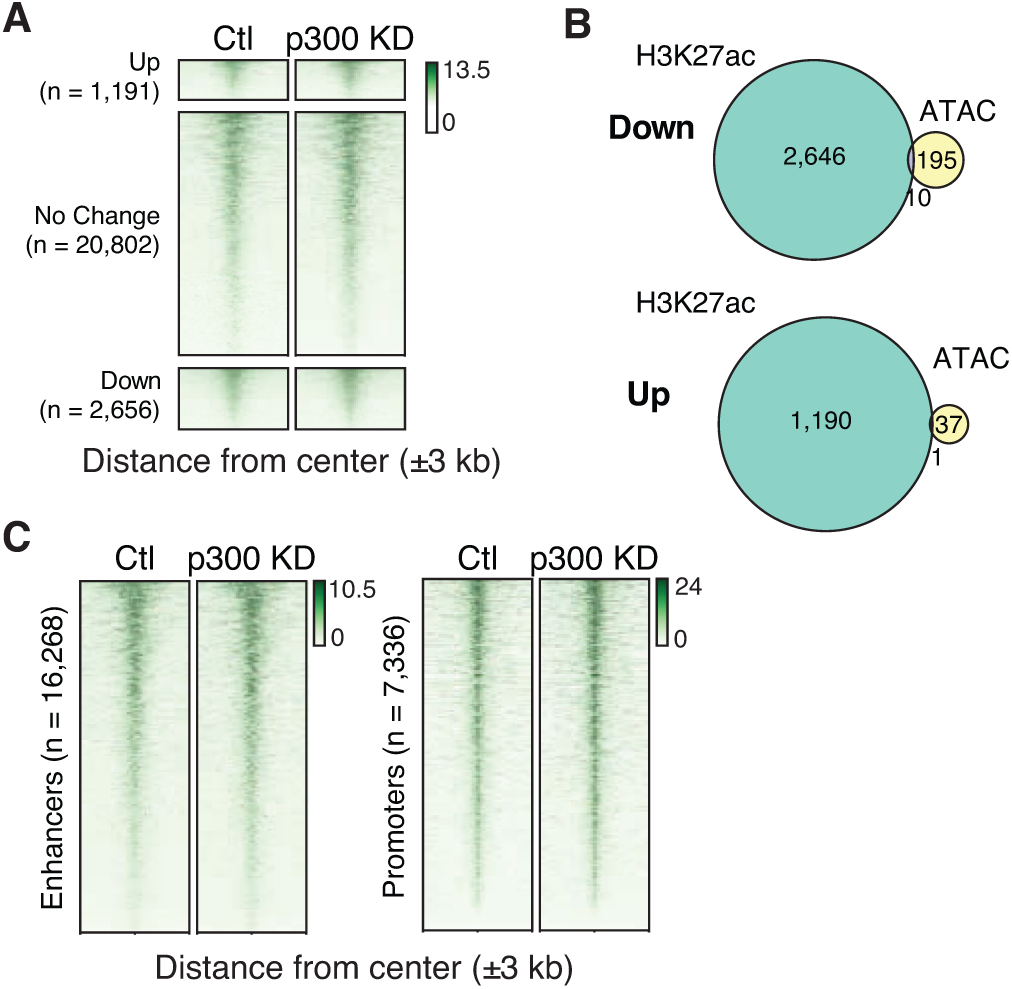
Chromatin Accessibility Is Independent of p300/CBP Levels in mESCs. (A) ChIP-seq heatmap of ATAC-seq in Ctl and p300 KD cells at regions that lose (down), maintain (no change), and gain (up) H3K27ac after p300 KD. Each row represents a single region. (B) Venn diagrams showing the relationship between ATAC-seq and ChIP-seq dysregulated regions after p300 KD. (C) ChIP-seq heatmap of H3K27ac at enhancers (left) and promoters (right) in Ctl and p300 KD mESCs.

**Suppl Figure 3:**
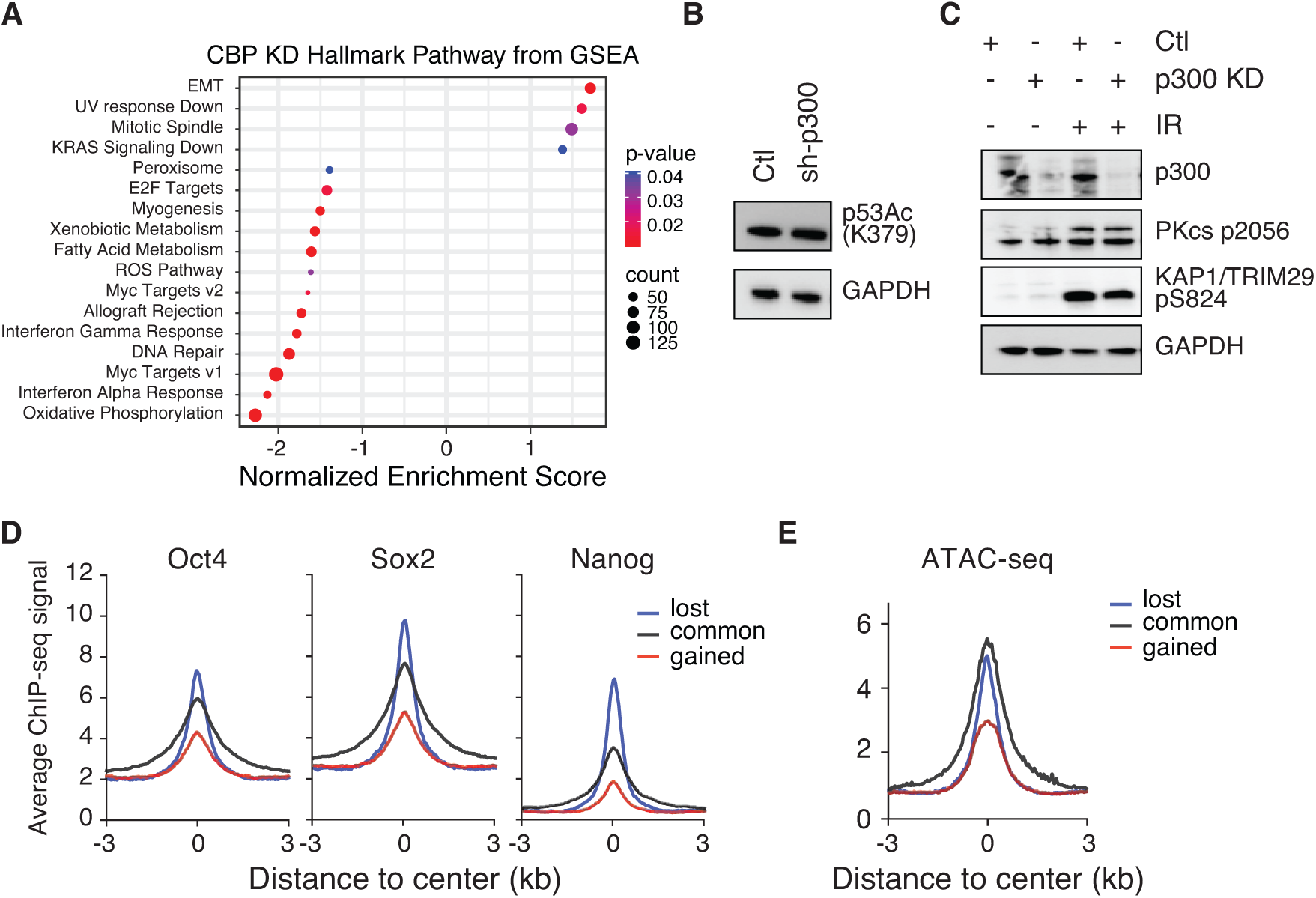
Loss of p300 leads to p53 increase. (A) GSEA pathway analysis of significantly regulated genes (based on RNA-seq) in CBP KD mESC compared to Ctl mESC. The normalized enrichment score (NES) is indicated for each gene set. (B) Immunoblot from whole cell lysates from mESCs transfected with scramble shRNA (Ctl) or p300 shRNA. (C) Immunoblot from whole cell lysates from mESCs transfected with scramble shRNA (Ctl) or p300 shRNA. Prior to harvesting, mESCs were treated for three min with 10 Gy of IR. (D) ChIP-seq average profiles of Oct4, Nanog, Sox2 in wild-type mESCs at regions of H3K27ac enrichment that are lost, common and gained after p300 KD. (E) ATAC-seq average profile in wild-type mESCs at regions of H3K27ac enrichment that are lost, common and gained after p300 KD.

